# A Conserved Metabolic–Oxidative Axis Underlies Immune Cell Cryo-vulnerability

**DOI:** 10.64898/2026.03.26.714376

**Authors:** Zujian Mo, Hongwei Yang, Meng Zhang, Huimei Cao, Lingqi Wang, Kun Tao, Xiaoshuai Chen, Caixia Han, Carlos Bustamante, Zhang Liu, Jianjun Wang

**Author notes:** Corresponding author: (J.W.); (Z.L.); (C.H.); (C.B.). Z.M., H.Y., and M.Z. contributed equally to this work.

## Abstract

Immunotherapy has emerged as a transformative approach for treating cancer and other diseases, yet its widespread deployment requires effective cryopreservation strategies to enable scalable global distribution. However, many immune cell types remain acutely vulnerable to freeze-thaw stress, and the underlying mechanisms of this cryo-vulnerability are not well defined. In this study, we combined metabolic flux analysis, ROS quantification, lipidomics, and preclinical xenograft models to investigate how metabolic state influences cryopreservation outcomes. We found that immune cell activation induces a metabolic shift characterized by elevated glucose utilization and excessive ROS production, leading to profound post-thaw loss of viability and function, as demonstrated by a ∼25% survival rate in natural killer (NK) cells. Targeted pretreatments—including inhibitors of glucose metabolism, antioxidants, and suppression of lipid peroxidation—restored post-thaw recovery to nearly 90% while preserving effector activity and antitumor efficacy. Similar protective effects were observed across αβ T cells, γδ T cells, and macrophages, defining a conserved metabolic–oxidative pathway of cryo-vulnerability and offering applicable strategies to enhance immune cell preservation.

## INTRODUCTION

The clinical success of adoptive cell therapies—particularly those utilizing genetically engineered immune cells—has reshaped the therapeutic landscape for hematologic malignancies and is expanding into solid tumors^1–3^. Despite these advances, the widespread clinical deployment of these sensitive, living medicines remains constrained by substantial logistical challenges across manufacturing, storage, and distribution^4–6^. Cryopreservation therefore represents a critical enabling technology, linking centralized production with global delivery while maintaining product potency. This need is especially acute for emerging allogeneic immune cell therapies^7–9^, whose scalable, off-the-shelf deployment requires cryopreservation approaches specifically tailored to immune cells.

Currently, therapeutic immune cell cryopreservation remains an evolving approach for long-term clinical use, with substantial challenges in preserving post-thaw immune function^10^. In contrast to conventional cell types, such as stem cells and established cell lines, which typically retain high viability and function following standard controlled-rate freezing with 10% DMSO^11–13^, immune cells often undergo delayed-onset death and marked functional impairment^14–21^. In T cells, cryopreservation induces apoptotic priming, marked by phosphatidylserine (PS) externalization, leading to reduced cytokine secretion, impaired proliferation, and mitochondrial dysfunction^14–16^. Natural killer (NK) cells display greater fragility, with viability falling from ∼95% to ∼20% after freezing and a complete loss of in vivo expansion potential along with reduced cytotoxicity, interferon-gamma (IFN-γ) production, and migratory capacity^17–19^. Antigen-presenting cells, such as dendritic cells (DCs), also exhibit functional impairment after freezing, including loss of lymph node homing and T-cell priming^20,21^. Collectively, these structural and functional deficits highlight that immune cells, unlike conventional cell types, undergo distinct cryo-induced injury that compromises their therapeutic utility, underscoring the urgent need for mechanistic insights to inform next-generation preservation strategies.

In this study, we demonstrate that the heightened metabolic activity of immune cells upon activation is associated with reduced resilience to cryopreservation. Activated NK cells, characterized by increased glucose metabolism, were observed to accumulate reactive oxygen species (ROS). Interventions that limit glucose metabolism, scavenge ROS, and enhance resistance to lipid peroxidation, which sequentially influence one another, significantly improved post-thaw cell viability and effector function, indicating their interconnected roles within a shared metabolic–oxidative axis that underlies cryosensitivity. Using preclinical HepG2 xenograft mouse models, we validated that NK cells pretreated with these interventions retained superior tumor control post-thaw compared to those cryopreserved by conventional methods. These results support a metabolism-driven ROS accumulation model of cryo-vulnerability in NK cells. Importantly, we found that this model was not restricted to NK cells but also extended to other immune cell types, including αβ T cells, γδ T cells, and macrophages, suggesting that pre-freeze metabolic activation and elevated ROS levels broadly contribute to cryoinjury. Together, these findings establish a unifying mechanistic framework for cryopreservation-induced damage across diverse immune cells and present a simple, applicable strategy to attenuate freezing-induced injury.

## RESULTS

### Differential cryotolerance between activated and non-Activated NK cells

Natural killer (NK) cells are widely used in cancer immunotherapy and are commonly activated ex vivo to enhance cytotoxic and cytokine-secreting functions prior to clinical application. Recent work has shown that cryopreservation-induced apoptosis in NK cells is driven by cytosolic leakage of granzyme B from destabilized cytotoxic granules, whereas pretreatment with IL-15 and IL-18 improves post-thaw recovery^18^. Because these cytokines are central regulators of NK cell activation^22^, this convergence suggests that cryo-vulnerability may reflect a state-dependent property governed by activation-linked upstream reprogramming, rather than an intrinsic limitation of NK cells. Such a framework could reconcile conflicting reports describing cryopreservation as either severely impairing^17–19^ or having limited effects on^23,24^ NK cell function. To date, however, direct side-by-side comparisons of activated and non-activated NK cells under identical cryopreservation conditions are lacking, leaving this principle untested.

To assess whether the activation state affects cryopreservation, we isolated NK cells (non-activated) and expanded them (activated) from healthy donors (Figure S1A), then cryopreserved both populations using a commercial cryoprotective solution (NK Cell Cryopreservation Media) according to established protocols (see Materials and Methods). Interestingly, the two populations exhibited significant differences in cryotolerance: AO/PI assay showed that activated NK cells displayed significantly higher post-thaw mortality compared to non-activated counterparts (Figure 1A), with apoptosis evident among the dying cells (Figure 1, B and C). Consistently, quantitative analysis showed that post-thaw recovery rates of activated NK cells declined to approximately 20% after 24 hours (Figure 1, D and E). To test functional integrity, we assessed the cytotoxicity of NK cells after cryopreservation against K562 target cells. After cryopreservation, activated NK cells showed significantly impaired function (Figure 1F). In contrast, non-activated NK cells demonstrated robust cryoresilience, with post-thaw recovery rates exceeding 90% and remaining stable over a 24-hour incubation (Figure 1, G and H). These cells also retained post-thaw activation and expansion capacity, with proliferation kinetics comparable to freshly processed controls (Figure S1B). Importantly, cryopreserved expanded NK cells derived from non-activated precursors exhibited cytotoxicity equivalent to their fresh counterparts (Figure 1I). Repeating above experiments with a GMP-compliant cryopreservation medium (CryoStore 10), widely used in clinical settings, also yielded similarly poor outcomes for activated NK cells (Figure S1C).

**Figure 1.**
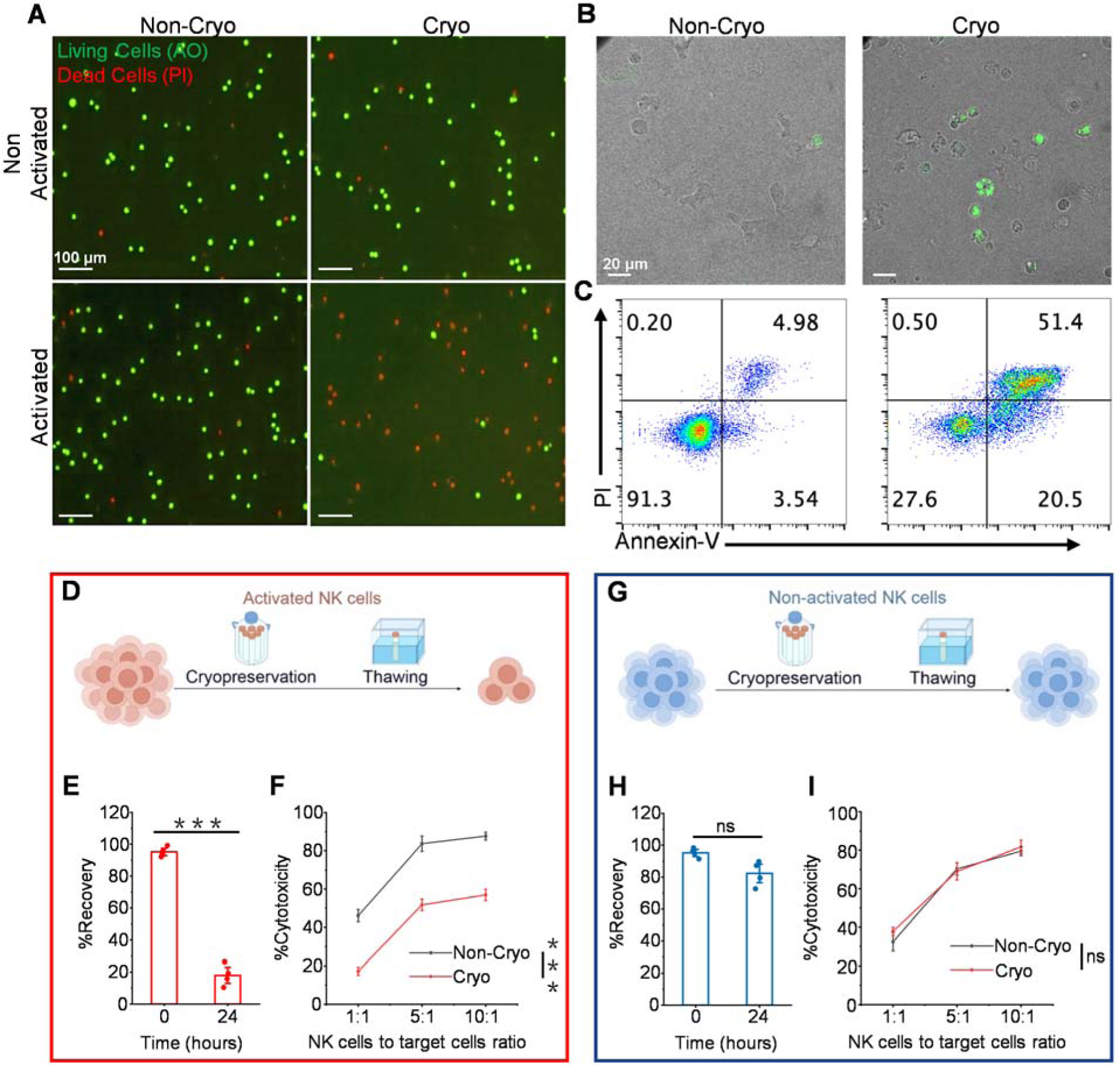
Discovery of differential cryotolerance between activated and non-activated NK cells. (A) Viability of NK cells detected by AO/PI dyes. Green channel (AO) indicates living cells and red channel (PI) indicates dead cells. (B) Confocal microscopy imaging showing the apoptosis cells (green channel) in fresh versus cryopreserved activated NK cells. (C) Apoptosis of activated NK cells analyzed by Annexin V/PI staining before and after cryopreservation. (D) A schematic demonstrating the cryopreservation tolerance of the Non-activated NK cells. (E) Non-activated NK cell recovery ratio at 0 and 24 h after thawing. The ratio defined as the viable cells after thawing relative to pre-freeze counts (n = 4 healthy donors). (F) Cytotoxicity of NK cells expanded from non-cryopreserved or cryopreserved non-activated NK cells at 1:1, 5:1 and 10:1 effector: target ratio for 4 h (n = 3 healthy donors). (G) Schematic demonstrating a substantial loss of cell numbers in activated NK cells following cryopreservation. (H) Activated NK cell recovery 0 and 24 h after thawing (n = 4 healthy donors). (I) Activated NK cytotoxicity assay with K562 cells with different effector to target ratio (n = 3 healthy donors). Data are the mean ± SEM (E, F, H, and I). Statistical analysis was performed using an unpaired two-tailed Student’s t test. Ns, P > 0.05, ***P < 0.001. Cryo: cryopreserved cells; Non-Cryo: non-cryopreserved cells.

These results demonstrate that activated NK cells exhibit markedly reduced post-cryopreservation recovery and cytotoxicity compared to their non-activated counterparts. This notable difference in cryotolerance between the two cellular states raises an intriguing question: what cellular mechanisms underlie the heightened susceptibility of activated NK cells to cryodamage?

### Inhibiting NK92 cell activation pathway improves resistance of NK92 cells to cryopreservation

To investigate the mechanisms underlying differential cryotolerance between activated and non-activated NK cells, we employed a reverse-engineering approach by inhibiting activation. For this, we selected the NK92 cell line as a model system, as it is a well-characterized, and it is a strictly Interleukin-2 (IL-2)–dependent NK cell line whose activation can be modulated using pharmacological interventions targeting signaling pathways downstream of the IL-2 receptor^25,26^. As validation, we show that cryopreservation of activated NK92 cells also resulted in a marked decline in both post-thaw recovery and cytotoxic function, closely mirroring the response observed in activated primary NK cells (Figure 2, A and B). This parallel behavior suggests that activation-associated cryo-vulnerability may be a generalizable phenomenon across NK cell systems.

**Figure 2.**
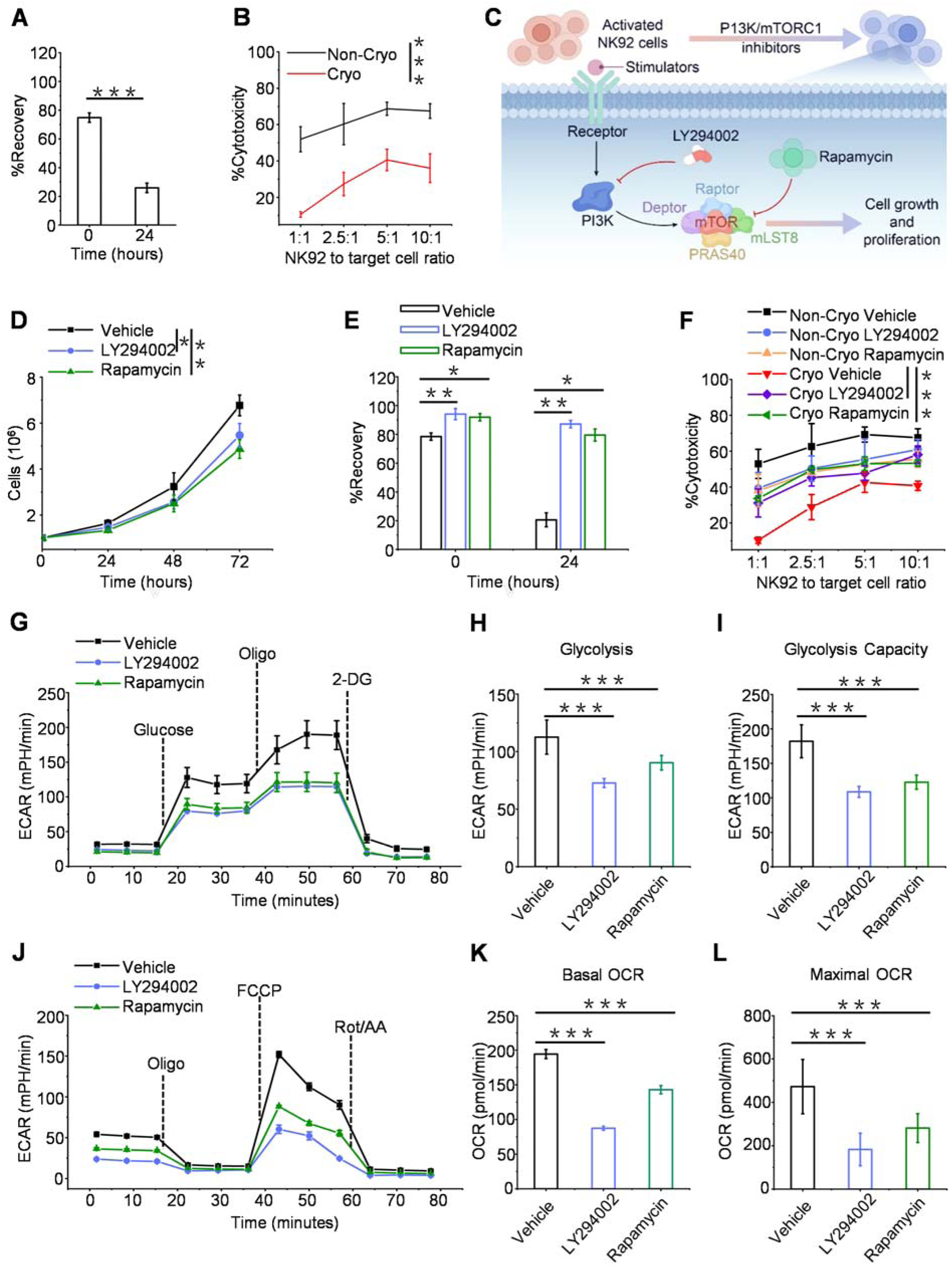
Inhibiting NK cell activation pathway improves resistance of NK cells to cryopreservation. (A) NK92 cell recovery at 0 and 24 h after thawing. (B) NK92 cytotoxicity assay before and after cryopreservation with Raji cells. NK92 cells were cultured with Raji cells at 1:1, 5:1 and 10:1 effector:target ratio for 4 h. (C) Schematic of the experimental workflow for PI3K/mTORC1 inhibition in NK92 cells using LY294002 (25 μM) and rapamycin (5 μM). (D) Proliferation kinetics of NK92 cells treated with LY294002 (25 μM) or rapamycin (5 μM). (E) Post-thaw recovery of inhibitor-pretreated NK92 cells (0 h and 24 h) relative to pre-cryopreservation viable cell counts. (F) LY294002 (25 μM) and Rapamycin (5 μM)-pretreated NK92 cytotoxicity assay with Raji cells. (G) Real-time extracellular acidification rate (ECAR) profiles of inhibitor-treated NK92 cells, measured by Seahorse XF analysis. (H) Glycolytic activity of inhibitor-treated NK92 cells. (I) Glycolytic capacity of inhibitor-treated NK92 cells. (J) Real-time oxygen consumption rate (OCR) profiles of inhibitor-treated NK92 cells. (K) Basal mitochondrial respiration in inhibitor-treated NK92 cells. (L) Maximal mitochondrial respiratory capacity in inhibitor-treated NK92 cells. Data are the mean ± SEM, n = 3 biologically independent experiments (A, B, D to F, H, I, K, and L). Statistical analysis was performed using an unpaired two-tailed Student’s t test. *P < 0.05, **P < 0.01, ***P < 0.001. Cryo: cryopreserved cells; Non-Cryo: non-cryopreserved cells.

Using the immunosuppressive agents LY294002 (PI3K inhibitor)^27^ or rapamycin (mTORC1 inhibitor)^28^ to suppress NK92 activation and proliferation (Figure 2, C and D), we observed significantly enhanced post-thaw recovery, with viability increasing from ∼25% to over 80% (Figure 2E). Moreover, the cytotoxic function of NK92 cells treated with LY294002 or rapamycin was largely preserved after cryopreservation, remaining comparable to pre-freeze levels (Figure 2F), indicating enhanced cryoresilience. These findings are consistent with a model in which PI3K/mTORC1 pathway activation contributes to cell cryo-vulnerability. However, it is noteworthy that the cytotoxicity of inhibitor-treated NK92 cells was modestly reduced (Figure 2F), possibly due to reduced expression of effector molecules, granzyme B (GZMB) and IFN-γ (Figure S2, A and B).

We note that the PI3K/mTORC1 signaling pathway mediates NK cell activation through metabolic reprogramming, which upregulates the expression of nutrient transporters and metabolic enzymes to meet the increased biosynthetic and bioenergetic demands of cell growth and proliferation^29,30^. To evaluate the metabolic consequences of PI3K/mTORC1 inhibition, we performed seahorse metabolic flux analysis. LY294002-and rapamycin-treated NK92 cells showed a significant reduction in both glycolysis (Figure 2, G to I) and oxidative phosphorylation (OXPHOS) (Figure 2, J to L)—two glucose-fueled metabolic pathways characteristic of the activated NK cell state^31^. These findings hint at a possible mechanistic link between elevated metabolic activity and susceptibility to cryopreservation, suggesting that the downstream metabolic reprogramming accompanying NK cell activation may contribute to freeze–thaw vulnerability.

### Reduced glucose metabolism and ROS production decrease NK92 cell cryo-vulnerability

Metabolic alterations are a hallmark of NK cell activation, enabling cells to meet the heightened biosynthetic and energetic demands of proliferation and cytokine production. As shown above, inhibition of PI3K/mTORC1 in activated NK cells downregulated both glycolysis and OXPHOS, which correlated with reduced cryo-sensitivity. These findings prompted us to directly investigate the roles of glucose metabolism and OXPHOS in contributing to NK cell cryo-vulnerability.

Using the fluorescent glucose analog 2-(NBD) Amino-2-deoxyglucose (2-NBDG) as a probe^32^, we quantified the glucose uptake in activated vs. activation-inhibited NK92 cells. Suppression of activation with LY294002 or rapamycin significantly reduced glucose transport (Figure 3A). To address whether attenuating glucose metabolism downstream of activation can directly improve cryoresistance, we pharmacologically inhibited glycolytic flux using 2-deoxy-D-glucose (2-DG), a synthetic glucose analogue that competitively inhibits hexokinase, thereby disrupting the Warburg-like metabolic phenotype characteristic of activated NK cells^33^. Pretreatment with 2-DG prior to cryopreservation significantly improved post-thaw recovery of NK92 cells, closely mirroring the effects observed with upstream PI3K/mTORC1 inhibition (Figure 3B). These results indicate that glucose metabolism is a critical downstream effector of activation that contributes to cryo-vulnerability in NK cells. Consistent with this model, 2-DG treatment also impaired cell proliferation and reduced expression of key effector molecules, including GZMB and IFN-γ (Figure S3, A to C), similar to the effects observed with PI3K/mTORC1 inhibitors.

**Figure 3.**
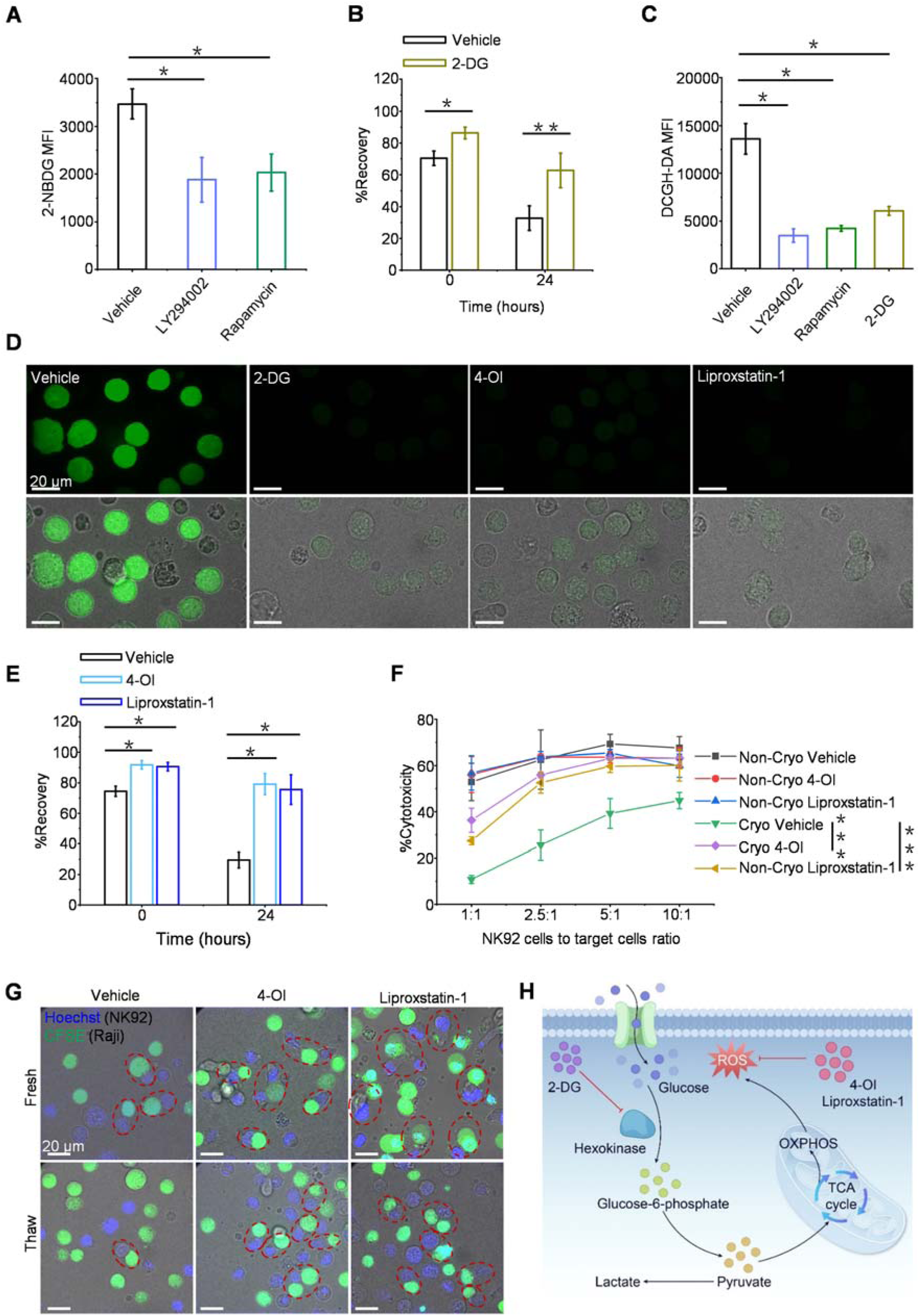
Glucose metabolism and ROS underlie NK92 cell cryo-vulnerability. (A) 2-NBDG uptake by LY294002 (25 μM) and Rapamycin (5 μM)-treated NK92. (B) 2-DG (5 mM)-pretreated NK92 cell recovery at 0 and 24 h post-thawing. (C) ROS levels of LY294002 (25 μM), Rapamycin (5 μM), and 2-DG (5 mM)-treated NK92 were detected by DCFH-DA probes. (D) Confocal microscopy imaging to assess the ROS levels in NK92 cells treated with vehicle or drugs was performed using DCFH-DA probes. (E) 4-OI (50 μM) and Liproxstatin-1(10 μM)-pretreated NK92 cell recovery 0 and 24 h after thawing. (F) 4-OI (50 μM) and Liproxstatin-1(10 μM)-pretreated NK92 cytotoxicity assay with Raji cells. (G) Confocal microscopy imaging showing the tumor-targeting adhesion capacity of NK92 cells [NK92 cells were stained with Hoechst (blue), tumor treated by CFSE (green) and cells adhering within the red dashed line]. (H) Glucose metabolism promotes the production of ROS. Data are the mean ± SEM, n=3 biologically independent experiments (A to C, E, and F). Statistical analysis was performed using an unpaired two-tailed Student’s t test. Ns, P > 0.05, *P < 0.05, **P < 0.01, ***P < 0.001. Cryo: cryopreserved cells; Non-Cryo: non-cryopreserved cells.

OXPHOS operates downstream of glycolysis and the TCA cycle, using NADH and FADH₂ to fuel the electron transport chain for ATP production, and is also the main cellular source of reactive oxygen species (ROS). Previous studies have established a strong correlation between enhanced glycolytic flux and excessive ROS generation^34,35^, which can impair cellular fitness^35^. To directly evaluate this link in NK92 cells, we measured intracellular ROS using DCFH-DA–based flow cytometry^37^. Both direct inhibition of glycolysis (2-DG) and upstream inhibition of PI3K/mTORC1 (LY294002 or rapamycin) significantly reduced ROS levels (Figure 3C), confirming that suppression of metabolic activation alleviates oxidative stress in NK92 cells.

To directly evaluate the role of ROS in cryodamage susceptibility, we pretreated NK92 cells with two antioxidants prior to cryopreservation: four-octyl itaconate (4-OI), which activates Nrf2 signaling to induce antioxidant enzyme expression^38^, and liproxstatin-1, a lipophilic ROS scavenger^39^. Both treatments notably reduced ROS levels comparable to 2-DG pretreatment (Figure 3D), and resulted in marked improvements in post-thaw recovery (Figure 3E). Importantly, NK92 cells pretreated with either antioxidant also demonstrated enhanced cytotoxic activity after thawing compared to untreated controls (Figure 3F). To further assess the preservation of effector functions, we employed confocal fluorescence imaging to visualize NK92-tumor cell interactions in co-culture. Cryopreserved NK92 cells exhibited significantly reduced tumor-targeting adhesion capacity (Figure 3G and Movie S1). In contrast, antioxidant-pretreated NK92 cells retained their ability to recognize and bind to tumor cells post-thaw, similar to cells that were not frozen, indicating preserved functional integrity.

Given that reactive oxygen species (ROS) are diffusible across cell membranes^40^, cells cultured at higher densities may experience elevated ROS levels due to intercellular transmission. We further investigated whether cell culture density impacts cryopreservation outcomes. NK92 cells cultured at low density exhibited significantly reduced ROS levels and enhanced post-thaw recovery compared to cells cultured at standard densities (Figure S3, D to F). These findings further support the idea that intracellular ROS accumulation contributes NK92 cell cryosensitivity. In summary, our results are consistent with a model in which elevated glucose metabolism, accompanied by excessive ROS production in activated NK92 cells, underlies their heightened susceptibility to cryodamage. Inhibiting glycolytic flux or directly scavenging ROS significantly improves cryopreservation outcomes (Figure 3H).

### Targeting lipid peroxidation prior to cryopreservation preserves NK92 cell integrity

The observed correlation between elevated ROS production and cryo-vulnerability in activated NK cells directed our attention to ROS as a potential mediator of cryodamage. ROS can injure cells by oxidizing multiple macromolecules, including proteins and nucleic acids. Among these none-specific targets, phospholipids are particularly susceptible due to their high content of polyunsaturated fatty acids (PUFAs), whose multiple double bonds are highly vulnerable to free radical attack^41^. ROS-induced lipid peroxidation disrupts membrane architecture by impairing lipid packing, reducing fluidity, and increasing osmotic fragility^42–45^. This vulnerability is especially critical during cryopreservation, when membrane integrity is further compromised by ice crystal formation, mechanical stress, and osmotic shock. These findings motivated us to quantitatively analyze lipid peroxidation level in NK92 cells and its impact on cryopreservation outcomes.

To assess NK92 cell lipid peroxidation levels, we employed BODIPY 581/591 C11, an oxidation-sensitive fluorescent probe that incorporates into membranes and shifts emission from red to green upon oxidation^46^. Validation experiments using NK92 cells pretreated with antioxidants 4-OI or liproxstatin-1 showed a marked reduction in BODIPY 581/591 C11 oxidation (Figure S4A), which suggests the alleviation of lipid peroxidation. While antioxidants target the ROS burden, lipid composition determines the substrate availability. To reduce the pool of ROS-susceptible lipid substrates, i.e. PUFAs, we pharmacologically induced the inhibition of ACSL4—an enzyme that facilitates the incorporation of PUFAs into NK92 cell membrane phospholipids, thereby, indirecrtly, promoting susceptibility to peroxidative damage^47^. NK92 cells pretreated with the ACSL4 inhibitors Rosiglitazone (Rosi)^47^ or AS252424^48^ displayed reduced levels of arachidonic acid (20:4)– and adrenic acid (22:4)–containing phosphatidylethanolamine (PE) species in lipidomics assay, confirming effective suppression of ACSL4 activity (Figure 4, A and B). In the meantime, the ACSL4 inhibition was accompanied by significantly decreased lipid peroxidation, as assessed by flow cytometer assays (Figure S4B) and BODIPY probes–based confocal imaging (Figure 4C). We noted that ACSL4-mediated suppression of lipid peroxidation demonstrated comparable efficacy than that of antioxidant treatments (Figure 4C).

**Figure 4.**
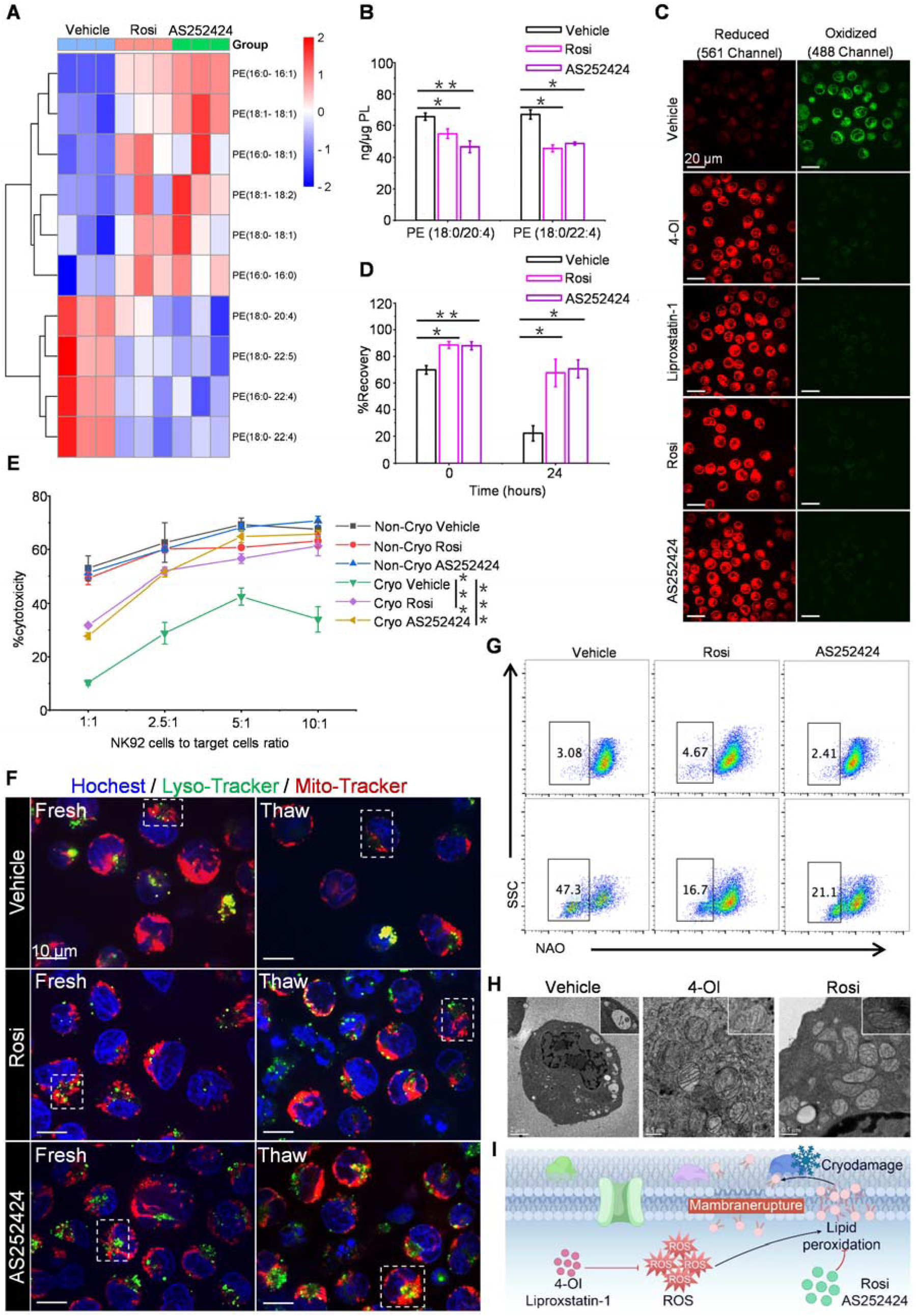
Targeting lipid peroxidation prior to cryopreservation preserves NK92 cell integrity. (A) Heatmap of major PE species in NK92 cells treated with Rosi (50 μM) or AS252424 (20 μM). Each PE species was normalized to the corresponding mean value. Each column represents an individual duplicate. The scale bar indicates Z-score. (B) Effects of Rosi (50 μM) or AS252424 (20 μM) on two major molecular species of PE (PE (18:0/20:4) and PE (18:0/22:4)) representing substrates for oxygenation. (C) Confocal microscopy imaging showing the lipid peroxidation levels of vehicle or pharmacologically treated NK92 cells using BODIPY dyes. (D) Rosi (50 μM) or AS252424 (20 μM)-pretreated NK92 cell recovery 0 and 24 h after thawing. (E) Rosi (50 μM) or AS252424 (20 μM)-pretreated NK92 cytotoxicity assay with Raji cells. (F) Confocal microscopy imaging showing the morphological characteristics of mitochondria (red) and lysosomes (green) in NK92 cells. Blue indicates nucleus. (G) Mitochondrial integrity of Rosi (50 μM) or AS252424 (20 μM)-pretreated NK92 was detected by NAO probes. NAOlo cells indicate cells with damaged mitochondia, NAOhi cells indicate cells with intact mitochondia. (H) Electron microscopy demonstrating subcellular structure of NK92 cells. Inset images: the morphological characteristics of mitochondria. Zooming into the structure of individual autophagosomes and mitochondria. (I) ROS promotes lipid peroxidation. Data are the mean ± SEM, n= 3 biologically independent experiments (B, D, E and G). Statistical analysis was performed using an unpaired two-tailed Student’s t test. Ns, P > 0.05, *P < 0.05, **P < 0.01, ***P < 0.001. Cryo indicates cryopreserved cells; Non-Cryo, non-cryopreserved cells.

To directly assess the role of lipid peroxidation in NK92 cell cryo-sensitivity, we measured post-thaw recovery and cytotoxicity as before. Pretreatment with Rosi or AS252424 markedly improved both parameters (Figure 4, D and E), underscoring the contribution of lipid peroxidation to cryodamage. Given previous reports that oxidative lipid damage compromises organelle fitness^49^, we next examined mitochondrial and lysosomal morphology changes in cryopreserved NK92 cells. Confocal microscopy revealed severe post-thaw disruption in mitochondrial and lysosomal integrity in vehicle-treated cells, with mitochondria shifting from elongated to fragmented morphology (red channel, dotted-line box) and lysosomes losing acid-responsive fluorescence (green channel, dotted-line box), whereas ACSL4 inhibitor–treated cells preserved both structures (Figure 4F). Nonyl acridine orange (NAO) is a specific fluorescent probe for mitochondrial integrity whose fluorescence signal weakens after mitochondrial membrane rupture^50^. NAO staining provided specific validation of mitochondrial damage post-thaw, which was attenuated in cryopreserved vehicle-treated cells but significantly mitigated by Rosi or AS252424 (Figure 4G). Similarly, lysosomal membrane permeability assessed with LysoTracker Red^51^ was elevated in cryopreserved cells but effectively reduced by ACSL4 inhibition (Figure S4C). Electron microscopy further confirmed these findings, revealing a large number of autophagosomes containing damaged mitochondrial in untreated cryopreserved NK92 cells, while those pretreated with ACSL4 inhibitors or antioxidants retained normal mitochondrial cristae structure^52^ (Figure 4H and inset).

Together, these results demonstrate that lipid peroxidation is one of the key downstream consequences of ROS and is directly linked to cryopreservation outcomes. Mitigating lipid peroxidation protects NK92 cells from cellular damage after cryopreservation, which might contribute to secondary cell death and impaired effector function after thawing. Targeting lipid peroxidation—either by reducing oxidative stress (antioxidant strategy) or limiting ROS-susceptible lipid incorporation (ACSL4 inhibition) (Figure 4I)—suggests a promising approach to preserve membrane integrity, thereby enhancing the recovery and functional competence of NK92 cells following cryopreservation.

### Translation of NK92 mechanistic findings to primary NK cells improves their antitumor efficacy in vivo

Our results in NK92 cells demonstrated that inhibition of key components along the glucose metabolism–ROS-lipid peroxidation pathway effectively alleviates cryo-vulnerability. To evaluate the transferability of these findings to primary NK cells, we first asked whether this hierarchical pathway is conserved, and then tested if inhibition of its critical nodes could similarly improve cryo-resilience. To this end, we downregulated the activation of expanded primary human NK cells using PI3K/mTORC1 inhibitors LY294002 and rapamycin. Consistent with our observations in the NK92 cell line, this treatment reduced glucose transport (Figure S5A). In addition, these PI3K/mTORC1 inhibitors, along with the glycolysis inhibitor 2-DG, decreased ROS production in activated NK cells (Figure S5B). We next pretreated activated NK cells with LY294002, rapamycin, 2-DG, the antioxidant 4-OI, or the ACSL4 inhibitor Rosi, all of which exert anti-lipid peroxidation effects (Figure 5A). Having confirmed the conservation of the pathway, we found that treatment with these drugs significantly enhanced the recovery of cryopreserved activated NK cells (Figure 5B). Moreover, cryopreserved NK cells preconditioned with either 2-DG or a combination of 4-OI and Rosi displayed cytotoxic activity against cancer cells similar to that of their fresh counterparts (Figure 5C).

**Figure 5.**
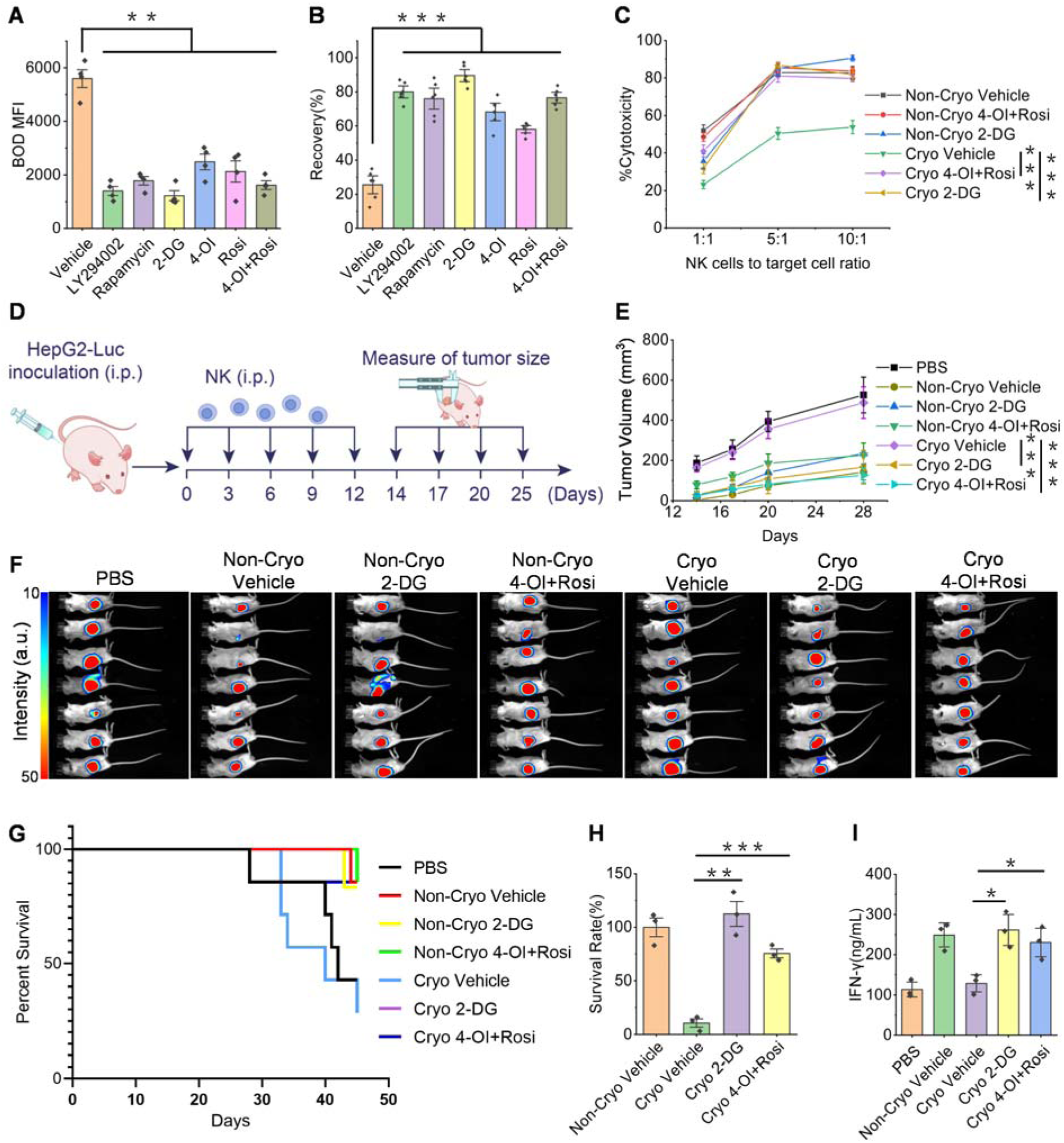
Mechanistic findings in NK92 cells translate to primary NK cells and improve antitumor efficacy in vivo. (A) BODIPY oxidation in LY294002 (25 μM), Rapamycin (2 μM), 2-DG (8 mM), 4-OI (50 μM) and/or Rosi (50 μM)-treated activated NK (n = 4 healthy donors). (B) LY294002 (25 μM), Rapamycin (2 μM), 2-DG (8 mM), 4-OI (50 μM) and/or Rosi (50 μM)-treated activated NK cell recovery 24 h after thawing (n = 4 healthy donors). (C) 2-DG (8 mM), or 4-OI (50 μM) + Rosi (50 μM)-pretreated activated NK cytotoxicity assay with K562 cells (n = 3 healthy donors). (D) Schematic diagram for in vivo experiment. (E) Tumor growth in mice (n = 7 mice/group). (F) Luminescent images of treated mice (n = 7 mice/group). (G) Survival curve of NK treated mice (n = 7 mice/group). The survival profile of the Cryo 2-DG cohort and Cryo 4-OI+Rosi cohort were overlapped by other groups in the Kaplan-Meier plot, with the 2-DG group exhibiting 100% survival rate,and Cryo 4-OI+Rosi group exhibiting 85.7% survival rate at the 45-day experimental terminus. (H) Normalized transplanted NK cell survival in mice (n = 3 mice/group). (I) IFN-γ levels of blood in mice (n = 3 mice/group). Data are the mean±SEM (A to C, E, H, and I). Statistical analysis was performed using an unpaired two-tailed Student’s t test. Ns, P >0.05, *P <0.05, **P <0.01, ***P <0.001. Cryo: cryopreserved cells; Non-Cryo: non-cryopreserved cells.

The true test of the cryostability of cell therapy products is how well their efficacy is retained after cryopreservation. To examine this aspect, we established a hepatoma carcinoma model using HepG2-Luciferase (Luc) cells, which are sensitive to NK cell-mediated killing. NOG mice were engrafted with both hepatocellular carcinoma and NK cells (Figure 5D). Cryopreserved NK cells pretreated with 2-DG or a combination of 4-OI and Rosi achieved significantly improved tumor control and prolonged survival of tumor-bearing mice compared to untreated cryopreserved NK cells (Figure 5E to G), indicating that pretreatment conserves NK cell potency during cryopreservation. In the disseminated xenograft lymphoma model using Raji cells, cryopreserved NK cells pretreated with 2-DG or a combination of 4-OI and Rosi similarly demonstrated enhanced tumor control capability compared to untreated controls (Figure S5, C to E). Consistent with these effects, pretreated NK cells also improved survival outcomes and IFN-γ secretion in peripheral blood relative to untreated cryopreserved NK cells, confirming that such pre-intervention treatments retain the activity of frozen NK cells (Figure 5, H and I).

### Generality of activation-induced cryovulnerability to immune cells

Our findings strongly support the hypothesis that activation-induced metabolic reprogramming, involving enhanced glucose utilization and ROS production, underlies the cryo-vulnerability of NK cells. Given that the activation of immune cells broadly drives metabolic upregulation and ROS accumulation^53–57^, we hypothesized that this phenomenon may not be unique to NK cells but instead represents a general feature of activated immune cells^14–21^. Accordingly, we sought to test this generality across additional immune cell types.

Following the same workflow as in NK92 cells, we treated expanded and activated primary human αβ T cells with LY294002 and rapamycin (Figure S6A) and observed a corresponding reduction in glucose transport (Figure S6B), consistent with our findings in activated NK cells. In addition, LY294002, rapamycin, and 2-DG treatment decreased ROS levels in activated αβ T cells (Figure S6C). To further investigate the impact of lipid peroxidation outcomes, activated αβ T cells were pretreated with LY294002, rapamycin, 2-DG, 4-OI, or Rosi, all of which exerted both anti-lipid peroxidative and cryoprotective effects on these cells (Figure 6, A and B, Figure S6D). Similar cryoprotective effects were observed in THP-1 cell-induced macrophages and expanded γδ T cells, suggesting the broad applicability of this preservation strategy across immune cell subtypes (Figure 6, C to E). Notably, in a hepatoma carcinoma model using HepG2-Luc cells, cryopreserved γδ T cells pretreated with 2-DG or a combination of 4-OI and Rosi allowed significantly improved survival, IFN-γ secretion, and tumor control compared to untreated cryopreserved γδ T cells in vivo (Figure 6, F to J). These findings confirm the cryostability of γδ T cell therapy products. Taken together, our findings support the conclusion that metabolism-induced ROS contributes to the susceptibility of activated immune cells to cryopreservation.

**Figure 6.**
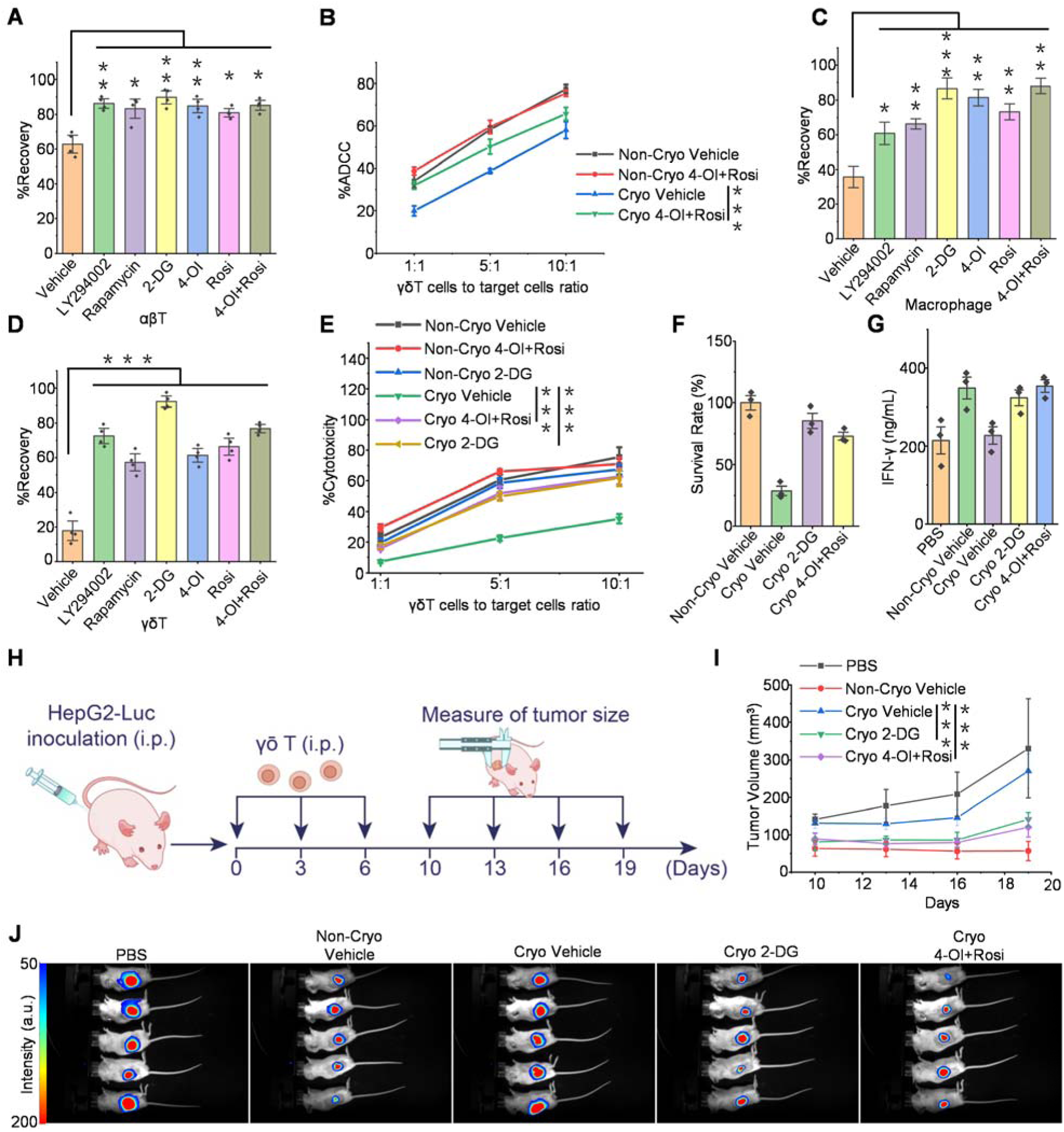
Generality of activation-induced vulnerability to cryopreservation. (A) LY294002 (25 μM), Rapamycin (2 μM), 2-DG (8 mM), 4-OI (50 μM) and/or Rosi (50 μM)-treated activated αβ T cell recovery 24 h after thawing (n = 4 healthy donors). (B) 4-OI (50 μM) and Rosi (50 μM)-pretreated activated αβ T cell ADCC assay using anti-CD3/CD20 antibody. T cells were cultured with Raji cells at the 5:1, 10:1 and 20:1 effector:target ratio for 24 h (n = 3 healthy donors). (C) LY294002 (25 μM), Rapamycin (2 μM), 2-DG (10 mM), 4-OI (200 μM) and/or Rosi (50 μM)-treated activated macrophages recovery 24 h after thawing. (D) LY294002 (25 μM), Rapamycin (2 μM), 2-DG (8 mM), 4-OI (50 μM) and/or Rosi (50 μM)-treated activated γδ T cell recovery 24 h after thawing (n = 4 healthy donors). (E) 2-DG (8 mM), or 4-OI (50 μM) + Rosi (50 μM)-pretreated activated γδ cytotoxicity assay with K562 cells (n = 3 healthy donors). (F) Normalized survival of mice transplanted γδ T cell (n = 3 mice/group). (G) IFN-γ levels of blood in treated mice (n = 3 mice/group). (H) Schematic diagram for vivo experiment. (I) Tumor growth in mice. (J) Luminescent images of treated mice (n = 5 mice/group). Data are the mean ± SEM (A to G, and I). Statistical analysis was performed using an unpaired two-tailed Student’s t test. Ns, P >0.05, *P <0.05, **P <0.01, ***P <0.001. Cryo: cryopreserved cells; Non-Cryo: non-cryopreserved cells.

## DISCUSSION

Immune cell therapy represents a promising advancement in immunotherapy, demonstrating significant efficacy against diverse cancers^1–3^. However, the lack of effective cryopreservation protocols necessitates labor-intensive and time-sensitive manufacturing immediately before treatment to achieve optimal therapeutic outcomes. This limitation not only creates critical challenges in clinical applications such as treatment delays, high costs, and poor accessibility, but also hinders the development of next-generation "off-the-shelf" and universal immunocyte therapies^4–6^. Overcoming technical barriers in immune cell cryopreservation is therefore essential to reduce costs, improve accessibility, and ultimately enabling their routine use as reliable, conventional treatments—shifting the field from "bespoke customization" to "accessible healthcare". NK cells, a central platform for immunotherapy, illustrate both the promise and the obstacles of cryopreservation. The basis of their cryosensitivity has not been fully defined, and reports have been contradictory—some describing little post-thaw effect on viability and function^23,24^, while others reporting severe impairments^17–19^. In this study, we identify activation status as the decisive outcome factor following cryopreservation. We find that activated NK cells are highly cryo-vulnerable, undergoing marked post-thaw death and loss of cytotoxicity, whereas non-activated NK cells display robust cryoresilience, maintaining high viability, proliferative potential, and potent cytotoxicity upon subsequent activation.

Because NK cell activation and expansion are integral to therapeutic preparation, understanding the factors underlying their reduced cryotolerance is important for improving preservation strategies. In our study, pretreatment of NK92 cells with PI3K/mTORC1 pathway inhibitors prior to cryopreservation significantly improved post-thaw recovery rates while maintaining cytotoxic function comparable to pre-freeze levels. The PI3K/mTORC1 signaling axis is a well-established regulator of immune cell activation, promoting the expression of nutrient transporters and metabolic enzymes required for effector function^29–31^. Consistent with this role, we observed that inhibition of the activated state through PI3K/mTORC1 signaling pathway indeed reduced glucose transport and glucose metabolism in NK92 cells. Interestingly, direct suppression of glycolysis with 2-deoxy-D-glucose (2-DG) prior to cryopreservation also significantly improved NK92 cryotolerance, with effects comparable to PI3K/mTORC1 inhibition. Therefore, these results indicate that activation-driven glucose metabolism significantly influences the cryosensitivity of NK92 cells (Figure 7, left).

**Figure 7.**
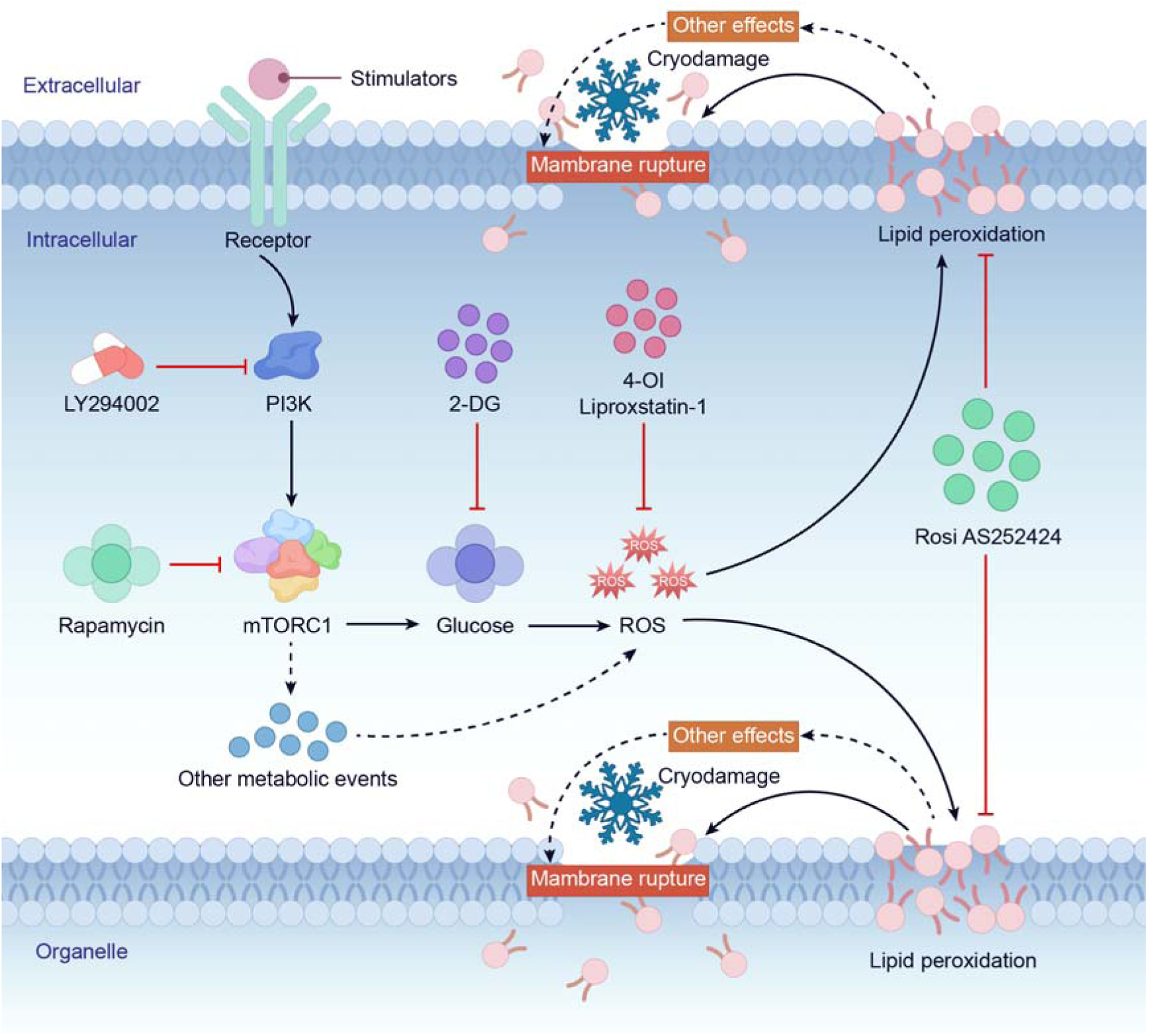
Proposed mechanisms of activation-induced cryosensitivity in immune cells. Solid lines denote established associations, while dashed lines represent potential associations.

Active glucose metabolism can also drive excessive reactive oxygen species (ROS) production^34,35^, which impairs cellular fitness^36^. Our data revealed that attenuating the activated state—either through inhibition of PI3K/mTORC1 signaling pathways or suppression of glucose metabolism—significantly reduced intracellular ROS levels in NK92 cells. Moreover, pretreatment with antioxidants to directly scavenge ROS before freezing further improved post-thaw recovery and cytotoxic function. Together, these findings support a model in which activation-induced glucose metabolism elevates ROS levels, thereby compromising the resistance of NK92 cells to cryodamage (Figure 7, middle). While glucose metabolism is the predominant source of ROS, we do not rule out additional pathways such as NADPH oxidase and glutamine metabolism that may also contribute to these species^58,59^.

ROS are known to damage a wide range of cellular biomolecules. Our findings identify phospholipids as a critical downstream target through which ROS mediate cryo-vulnerability (Figure 7, right). The high content of polyunsaturated fatty acids (PUFAs) renders membranes especially prone to peroxidation^41^, disrupting bilayer packing, reducing fluidity, and weakening osmotic resilience during freeze–thaw stress^42–45^. By showing that ACSL4 inhibition reduces PUFA incorporation, suppresses lipid peroxidation, and preserves membrane integrity, we establish lipid peroxidation as a direct driver of cryodamage rather than a byproduct of oxidative stress. Moreover, lipid peroxides can propagate damage to proteins and nucleic acids, providing a mechanistic explanation for the impaired survival and function of activated NK cells after thawing. These results highlight lipid peroxidation as a critical downstream consequence of ROS during cryopreservation and support ACSL4 inhibition as a mechanism-based strategy, complementary to antioxidant approaches, to enhance the stability of therapeutic NK cells.

Fragility during cryopreservation has been widely reported across immune cells—including NK cells, T cells, and dendritic cells^14–21^. Using NK92 cells as a prototype, we established an activation-driven metabolic cryo-vulnerability model in which fragility is linked downstream of elevated glycolytic flux, ROS overproduction, and lipid peroxidation (Figure 7). In this model, we do not rule out contributions from other metabolic pathways to ROS accumulation or alternative ROS targets beyond lipids. However, inhibition of any individual node in the pathway significantly improved cryopreservation outcomes, suggesting that activation-induced oxidative stress constitutes a convergent driver of cryo-vulnerability. Interestingly, this model can be generalized to primary NK cells as well as other immune cell types, including αβ T cells, γδ T cells, and macrophages. Pre-cryopreservation inhibition of glucose metabolism or alleviation of oxidative stress substantially reduced cryopreservation-induced damage in the aforementioned immune cells and preserved functionality in vitro and in vivo at levels comparable to their pre-freezing state. Collectively, these findings provide mechanistic insights into cryopreservation-induced damage in immune cells and establish a mechanism-guided preservation framework to enhance the stability of immune cell–based therapeutics. By addressing the critical barrier of long-term storage, this work is expected to accelerate the development and clinical translation of next-generation immune cell therapies.

Indeed, enhanced metabolism and elevated ROS levels represent a commonly observed phenomenon that extends beyond immune cells to diverse cell types and physiological systems. For instance, pancreatic islet cells express high levels of glucose-metabolizing enzymes for blood glucose sensing and exhibit substantial cell death and functional impairment post-cryopreservation, which significantly limits their clinical application in transplantation therapy for diabetes mellitus^60–63^. Cancer cells often display robust metabolism and high ROS levels^64^. Cells subjected to environmental stress, such as ischemia-reperfusion^65^, radiation^66^, and aging^67^, also experience ROS-related challenges. Future studies should systematically evaluate whether metabolism or ROS inhibition can provide broad cryoprotection across diverse cell types. Moreover, the precise mechanisms through which ROS mediates cryopreservation sensitivity remain incompletely defined. Elucidating how ROS drives cryopreservation failure will enable next-generation preservation technologies that strategically target these deleterious pathways.

### Limitations of the study

The present investigation is primarily reliant on specific pharmacological interventions. To achieve more precise mechanistic characterization and targeted manipulation, future work should employ advanced methodologies such as genetic engineering, multi-omics analyses (e.g., transcriptomics, proteomics), or other refined techniques. These approaches are essential for generating more robust and definitive data and conclusions. Furthermore, the findings derived from this study necessitate further validation across a broader spectrum of cell types and tissues to establish their generalizability and physiological relevance.

## RESOURCE AVAILABILITY

### Lead Contact

Further information and requests for resources and reagents should be directed to and will be fulfilled by the Lead Contact Jianjun Wang.

### Materials Availability

This study did not generate new unique reagents.

### Data and Code Availability

All data produced for this manuscript are available at Zenodo (https://doi.org/10.5281/zenodo.18252487). This paper does not report original code. Any additional information required to reanalyze the data reported in this paper is available from the corresponding author upon request.

## AUTHOR CONTRIBUTIONS

Conceptualization: Z.M., Z.L., J.W.

Methodology: Z.M., Y.H., Z.L., J.W.

Investigation: C.H., Z.L., J.W.

Visualization: H.C., L.W.

Supervision: Z.L., J.W.

Writing—original draft: Z.M., Y.H.

Writing—review & editing: C.H., Z.L., J.W., M.Z., C.B.

Funding acquisition: Z.L., J.W.

## List of Supplementary Materials

Figures. S1 to S6

Movie S1

## METHODS

### Study design

The objective of this investigation was to elucidate the molecular mechanisms reponsible for cryopreservation-induced cellular damage in immune cells and to develop optimized preservation protocols. Utilizing the NK92 cell line as our principal experimental model, we systematically identified critical determinants of cryopreservation efficiency and formulated evidence-based intervention approaches. To comprehensively characterize immune cell functionality throughout the cryopreservation process, we implemented an integrated analytical platform incorporating flow cytometric analysis, confocal laser microscopy, transmission electron microscopy (TEM), and In vivo imaging systems. This multimodal assessment enabled precise quantification of essential biological parameters including post-thaw viability, cytokine secretion dynamics, and antitumor cytotoxic capacity under both in vitro and in vivo conditions. Detailed experimental protocols, including specific biological replicate numbers and statistical analysis methods, are provided in the respective figure legends and Supplementary Materials. All animal studies were conducted in strict compliance with institutional ethical guidelines for laboratory animal care and use. To minimize experimental bias, mice were randomly allocated to control or treatment groups by independent investigators, with group assignments based on standardized tumor volume measurements prior to intervention.

### Chemicals

AS252424 (T6208) and Rosi (T0334) were purchased from Topscience. Liproxstatin-1 (HY-12726), LY294002 (HY-10108), Rapamycin (HY-10219), 4-OI ( HY-112675), 2-NBDG (HY-116215), 2-DG (HY-13966), LysoTracker Red (HY-D1300), Zoledronic Acid (HY-13777), DCFH-DA (HY-D0940), D-Luciferin (HY-12591A), and BODIPY C11 ( HY-D1301) were purchased from MedChemExpress. NAO (A1372) was purchased from ThermoFisher.

### Cells lines

THP-1, Raji, Raji-Luc, K562, and K562-mbIL15-41BBL feeder cells were cultured in RPMI 1640 (Thermo Fisher, #11875085) supplemented with 10% fetal bovine serum (FBS) (Seradigm, #97068-086), and 1% penicillin–streptomycin (P/S) (Gibco, #15140122). HepG2-Luc cells were cultured in DMEM (Thermo Fisher, #11965092) supplemented with 10% fetal bovine serum (FBS) (Seradigm, #97068-086), and 1% penicillin–streptomycin (P/S) (Gibco, #15140122). NK92 cells were cultured in a commercial NK92 medium (Sunncell, SNLM-405), which was composed of MEMα+12.5% FBS+12.5% horse serum+0.2 mM Inositol+0.1 mM β-mercaptoethanol+ 0.02 mM Folic Acid+200 IU/mL recombinant IL-2+1% P/S. All cell lines in our laboratory were cultured at 37°C in humidified incubator with 5% CO_2_ and verified by STR profiling.

## METHOD DETAILS

### Primary human NK cell isolation and expansion

Human peripheral blood mononuclear cells (PBMCs) were extracted from whole blood of healthy adult donors by Ficoll (Sigma-Aldrich, F4375), using density gradient centrifugation. We have obtained informed consent from all participants. The non-activated NK was isolated using NK Cell Isolation Kit (Miltenyi, #130-092-657). As for the activated NK, PBMCs were stimulated by irradiated K562-mbIL15-41BBL as feeder cells, 200 IU/mL IL-2 (Peprotech, #200-02) and expanded for 15-20 days in X-VIVO 15 (Lonza, #02-060Q).

### Αβ T-cell activation and expansion

Human αβ T cells were isolated and activated from human peripheral blood mononuclear cells (PBMCs) using the T Cell TransAct™ kit (Miltenyi Biotec, #130-111-160), in strict accordance with the manufacturer’s protocol. The activated cells were then expanded ex vivo for a period of 10 to 15 days. The expansion culture was maintained in X-VIVO 15 serum-free medium (Lonza, #02-060Q) supplemented with 100 IU/mL of recombinant human IL-2 (Peprotech, #200-02) to support T cell growth and proliferation.

### γδ T-cell activation and expansion

γδ T cells were expanded from human PBMCs by initial activation with 5 μM Zoledronic Acid. The activated cells were then maintained in culture for a period of 10 to 15 days in X-VIVO 15 serum-free medium (Lonza, #02-060Q), which was further supplemented with 200 IU/mL recombinant human IL-2 (Peprotech, #200-02) to promote sustained proliferation and cell growth.

### Macrophage differentiation and stimulation

Macrophage differentiation was induced in THP-1 monocytes by 6-hour treatment with 100 ng/mL phorbol 12-myristate 13-acetate (PMA). After two washes with PBS to remove PMA, the differentiated macrophage-like cells were cultured for 24 hours in PMA-free medium. Subsequently, cells were stimulated with 100 ng/mL lipopolysaccharide (LPS) for 6 hours.

### Cell viability assay

Cell viability was measured using a cell counter (Nexcelom, Cellometer K2) according to the manufacturer’s instructions. Briefly, after being incubated with the AO/PI dyes, dead cells and living cells were determined and counted by detecting cell fluorescence.

### Cell cryopreservation and thawing

The cells were resuspended and cultured in fresh medium to achieve a final density of 5× 10^5^ cells/mL. LY294002, Rapamycin, 2-DG, 4-OI, Liproxstatin-1, Rosi, or AS252424 were added 48 h before cryopreservation either separately or in combination as indicated in the main text. 1 × 10^7^ cells were washed with medium then resuspended in 1 mL of NK Cell Cryopreservation Media (CELLSTORE, #CS-NK-D1) or CryoStor® CS10 (STEMCELL, #07930) before being frozen to −80°C in a cell freezer (Corning®,#432001) overnight, and then being transferred to the liquid nitrogen for long-term storage. Cryopreserved cells were thawed in a 37°C water bath.

### Cytotoxicity assay

The cytotoxicity of NK92 cells was evaluated using Raji cells as targets. Briefly, target cells were labeled with CFSE and co-cultured with effector cells at varying effector-to-target (E:T) ratios. Following a 4-hour incubation at 37°C, the cells were stained with 7-AAD for 15 minutes and analyzed by flow cytometry. Cytotoxicity was determined by the percentage of CFSE^+^ cells that were also 7-AAD^+^. The same protocol was adopted for assessing the cytotoxicity of primary NK cells, αβ T cells, and γδ T cells, except that K562 cells served as the target cells. For the αβ T cell ADCC assay, 2 ng/mL of bispecific [CD3×CD20] biobodies (Bio X Cell #SIM0008) were added to the co-culture.

### Apoptosis assay

Apoptosis in NK cells was assessed 24 hours following post-thaw resuscitation via flow cytometric analysis. An Annexin V-FITC/PI Apoptosis Detection Kit (Beyotime, #C1062S) was employed for this purpose, and the assay was conducted in full compliance with the manufacturer’s protocol. Specifically, approximately 1×10^5^ cells were collected, washed, and resuspended in 200 μL of Binding Buffer. The cell suspension was then subjected to dual staining with Annexin V-FITC and PI for 15 minutes under light-protected conditions. Apoptotic rates were determined immediately after the incubation period.

### Immunofluorescence assay

Immunofluorescence assays were performed with fluorophore-conjugated antibodies and flow cytometry. By staining with antibodies against the CD3 (BioLegend, #300439) and CD56 (BioLegend, #362518), NK cells were defined as CD3^−^CD56^+^ cells, and T cells were defined as CD3^+^CD56^−^ cells. After treating NK92 cells with LY294002, Rapamycin, or 2-DG for 24 hours, the expression of GZMB or IFN-γ in NK92 cells was detected by staining with the Cytofix/Cytoperm Plus Kit (BD, #554714) and antibodies against GZMB (Invitrogen, MA5-23639) or IFN-γ (Invitrogen, MHCIFG29).

### Glucose uptake analysis

Prior to the glucose uptake assay, cells (including NK92, primary NK cells, and T cells) were pretreated with LY294002 or Rapamycin for 24 hours. The cells were then exposed to 10 μM 2-NBDG for 8 hours at 37°C. The level of 2-NBDG incorporation, reflecting glucose uptake, was quantified by flow cytometry with the excitation and emission wavelengths set at 488 nm and 525 nm, respectively.

### Seahorse Extracellular flux analysis

Seahorse analysis experiments were performed according to the manufacturer’s protocol on a XF-96 Extracellular Flux Analyzer (Seahorse Bioscience). In brief, 24 h after treatment with LY294002 or Rapamycin, NK92 cells were seeded at a density of 1 × 10^5^/well in plates coated with poly-D-lysine (Beyotime, #C0313) in XF medium (Agilent, #102353-100). Then the ECAR and OCR were measured in real time using Seahorse XF Glycolysis Stress Test Kit (Agilent, #103020-100) and Seahorse XF Cell Mito Stress Test Kit (Agilent, #103015-100) respectively.

### ROS detection

Prior to ROS measurement, the cells (NK92, primary NK, or T cells) were cultured in fresh media and treated with various compounds (LY294002, Rapamycin, 2-DG, 4-OI, or Liproxstatin-1) for 48 hours. For the determination of intracellular ROS, the pretreated cells were incubated with 10 μM of the fluorescent probe DCFH-DA for 30 minutes at 37°C. Flow cytometric analysis was then conducted with the excitation and emission wavelengths set at 488 nm and 525 nm, respectively, to detect the oxidized DCF signal.

### Lipid peroxidation detection

After being cultured in fresh media, NK92, NK or T cells were treated with LY294002, Rapamycin, 2-DG, 4-OI, Liproxstatin-1, Rosi, or AS252424 either separately or in combination for 48 h. Then these cells were incubated with 5 μM BODIPY for additional 30 min at 37°C to measure cellular lipid peroxidation by flow cytometry using excitation and emission wavelengths of 488 and 525 nm.

### Lipidomic analysis

Seventy-two hours after being treated with Rosi or AS252424, lipids were extracted from approximately 1×10^7^ NK92 cells using the Bligh and Dyer method. The organic phase was dried and dissolved in 50 μL of mobile phase A [trichloromethane/isopropanol 2:1 (v/v)]. Then, the lipids were mixed with mobile phase B [acetonitrile/isopropanol/water 7:9:4 (v/v/v) and 10 mM ammonium acetate], and MS analysis of lipids was performed on a Q-Exactive hybrid quadrupoleorbitrap mass spectrometer (Thermo Fisher Scientific).

### Mitochondrial integrity analysis

Assessment of mitochondrial integrity in NK92 cells was performed via flow cytometric quantification of NAO fluorescence. This assay is predicated on the specific and high-affinity binding of NAO to cardiolipin, a phospholipid predominantly located in the inner mitochondrial membrane, wherein the fluorescence yield is directly proportional to the mitochondrial mass and membrane integrity. In this experiment, harvested NK92 cells were methodically stained with 5 μM NAO for a duration of 30 minutes at 37°C, with precautions taken to minimize light exposure. Post-staining, the cells were subjected to two washes with cold PBS to ensure the removal of any non-specific dye and were subsequently analyzed. The fluorescent signal was definitively measured using a flow cytometer configured with a 488 nm laser for excitation and a 525/30 nm bandpass filter for emission collection, providing a quantitative measure of mitochondrial status.

### Lysosome membrane permeability analysis

To evaluate lysosomal membrane permeability, NK92 cells were harvested and incubated with 100 nM LysoTracker Red for 30 minutes at 37°C under light-protected conditions. Following incubation, cells were washed twice with PBS to remove excess dye and resuspended in complete medium for immediate analysis by flow cytometry. The fluorescence intensity of LysoTracker Red, which accumulates in acidic compartments and serves as an indicator of lysosomal integrity, was quantified using excitation at 488 nm and emission at 585 nm.

### Confocal imaging and TEM imaging

All fluorescence labeling images were acquired by CSU-W1 spinning-disk confocal scanner unit (YOKOGAWA) (λex=405, 488, 561 and 640 nm. 100X oil objective lens and bandpass filter (460/50, 525/50, 600/52, 708/75) were used for collecting fluorescence, respectively. All the images were processed and analyzed using Image J software subsequently.

For ROS imaging, we used the DCFH-DA probe. NK92 cells treated with different conditions were co-incubated with DCFH-DA for 30 minutes, then the NK92 cells were washed with PBS and imaged subsequently. Excitation was performed using a 488 nm laser, and fluorescence was collected using a 525/50 bandpass filter, and all the images were presented at the same contrast.

For the Raji cells killing assay of NK92 cells, we stained NK92 cells with Hoechst 33258 dye and Raji cells with CFSE dye. Subsequently, we mixed NK92 cells with Raji cells at appropriate concentrations and seeded them into confocal dishes. After incubating in a cell culture incubator for 3 hours, we performed confocal dual-color imaging, using the 405 nm channel (460/50) to image the Hoechst dye and the 488 nm channel (525/50) to image the CFSE dye.

For the imaging of lipid peroxidation of NK92 cells, we used the BODIPY 581/591 C11 probe. After co-incubating with the NK92 cells for 30 minutes, confocal imaging was performed. The fluorescence of the oxidized and reduced forms of probes was collected using the 488 (525/50) and 561 nm (600/52) channels, respectively.

For the multicolor imaging of subcellular organelles in NK92 cells, we used the Hoechst 33258, lyso-Tracker red and mito-tracker deep red dye stained the nucleus, lysosomes, and mitochondria of NK92 cells according to standard staining procedures, respectively. In confocal microscopy, we used the 405 (460/50), 561 (600/52), and 640 nm (708/75) channels to image nucleus, lysosomes and mitochondria, respectively. The subsequent merging of images was completed using ImageJ software.

Preparation of biological samples and TEM imaging were performed and acquired from the laboratory of Prof. Wanzhong He, National Institute of Biological Sciences (NIBS), Beijing, China.

### Adoptive NK cell transfer

For the HepG2 xenograft model, NSG mice received intraperitoneal (i.p.) injections of 2×10 luciferase-expressing HepG2 cells (HepG2-Luc) and 2×10 NK cells simultaneously at Day 0, followed by additional NK cell injections (2×10 cells/mouse) on Days 3, 6, 9, and 12. Tumor progression was monitored by measurement of tumor volume Days 14, 17, 20, and 25 and bioluminescence imaging (IVIS Spectrum Imaging System) at Days 25 following i.p. administration of luciferin (150 mg/kg), with data analysis performed using Living Image Software. In the Raji lymphoma model, mice were inoculated with 5×10 Raji-Luc cells via tail vein injection and randomized at Day 3 post-inoculation. Treatment groups received 5×10 NK cells on Days 3, 7, and 10 post-tumor injection. Bioluminescence imaging was performed every 3-4 days starting from Day 0 post-NK cell administration. Peripheral blood collection via orbital bleeding was conducted at Day 3 post-treatment for analysis of NK cell survival and IFN-γ production in vivo. The survival rate of transplanted NK cells in mice was determined by flow cytometry using anti-human CD56 antibody (BioLegend, #362518) staining, with the survival rate of fresh NK cells set as 100%. The IFN-γ produced by human NK cells in mice was measured using an ELISA kit (Solarbio, #SEKH-0046) according to the manufacturer’s instructions. Control groups in both models received tumor cells alone with PBS vehicle.

### Adoptive γδ T cell transfer

NSG mice were inoculated intraperitoneally (i.p.) with 4×10 luciferase-expressing HepG2 cells (HepG2-Luc) and 2×10 γδ T cells simultaneously on Day 0, followed by booster injections of γδ T cells (2×10 cells/mouse) on Days 3 and 6. Tumor growth was assessed through serial measurements of tumor volume on Days 10, 13, 16, and 19, and by bioluminescence imaging using an IVIS Spectrum Imaging System on Day 20 after i.p. administration of luciferin (150 mg/kg). Image analysis was performed using Living Image Software. For evaluation of γδ T cell survival and IFN-γ production in vivo, peripheral blood was collected via orbital bleeding at Day 3 post-treatment (following tail vein injection of 5×10 cells/mouse). The survival rate of transplanted γδ T cells in mice was determined by flow cytometry using anti-human CD3 antibody (BioLegend, #300439) staining, with the survival rate of fresh γδ T cells set as 100%. The IFN-γ produced by transplanted γδ T cells in mice was measured using an ELISA kit (Solarbio, #SEKH-0046) according to the manufacturer’s instructions. Control groups received tumor cells alone with PBS vehicle in both experimental models.

### Statistical analysis

Data were presented based on at least three independent experiments as mean ± SEM. The comparison of indicated two groups was performed using Student’s t - test (two - tailed, unpaired). Statistical significance was denoted as *P < 0.05, **P < 0.01, ***P < 0.001, and non - significant results were labeled as "ns". All statistical details are provided in the figure legends.

## Supplementary Materials

**Figure S1.**
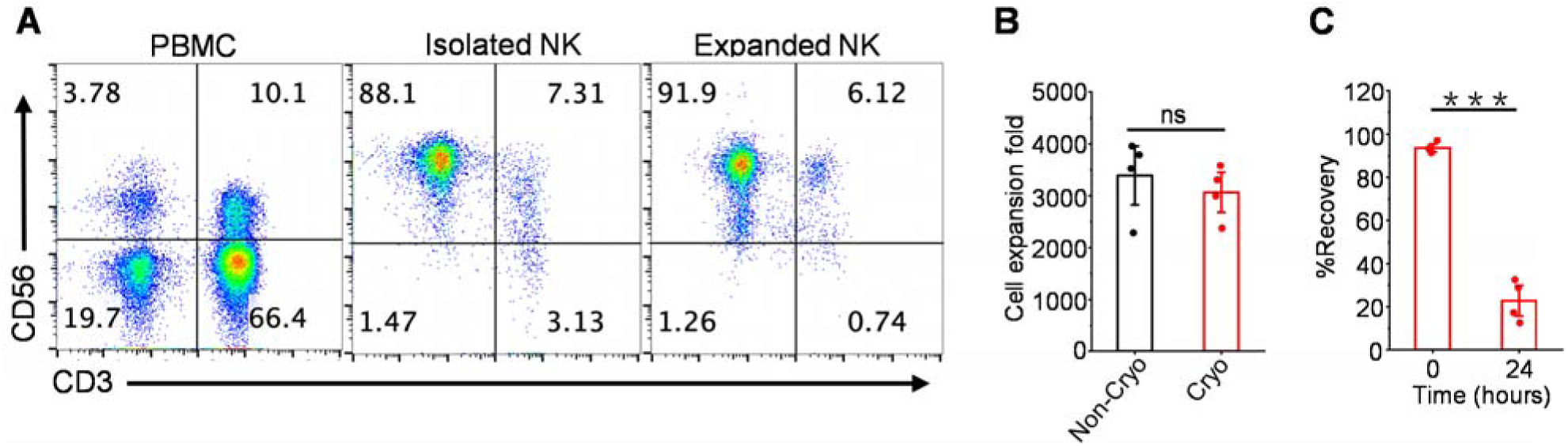
Characterizing NK cell phenotype, recovery, and proliferation. (A) Representative flow cytometry plots showing CD3^−^CD56^+^ NK cells. (B) Total fold expansion of peripheral blood derived NK cells 14 days after activation (n = 4 healthy donors). (C) Recovery of activated NK cells cryopreserved using CryoStor® CS10 at 0 and 24 h after thawing (n = 4 healthy donors). Data are the mean ± SEM (B and C). Statistical analysis was performed using an unpaired two-tailed Student’s t test. Ns, P > 0.05, ***P < 0.001. Cryo: cryopreserved cells; Non-Cryo: non-cryopreserved cells.

**Figure S2.**
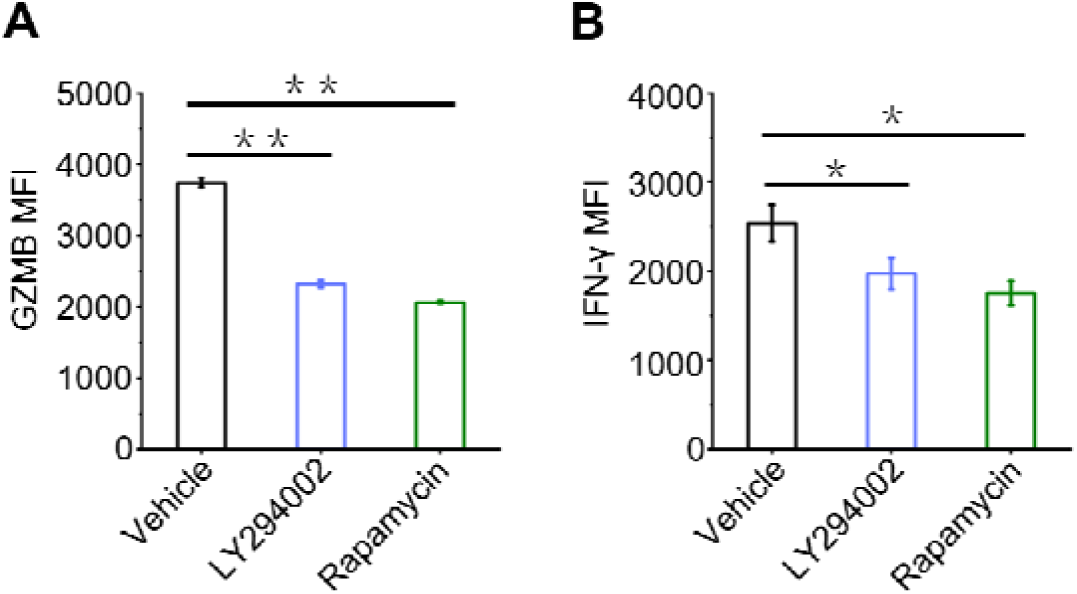
Effects of LY294002/Rapamycin on the expression of cytokines of NK92 cells. (A) GZMB levels in NK92 cells treated with LY294002 (25 μM) or Rapamycin (5 μM). (B) IFN-γ levels of LY294002 (25 μM) or Rapamycin (5 μM)-treated NK92 cells. Data are the mean ± SEM, n = 3 biologically independent experiments (A and B). Statistical analysis was performed using an unpaired two-tailed Student’s t test. *P < 0.05, **P < 0.01, ***P < 0.001.

**Figure S3.**
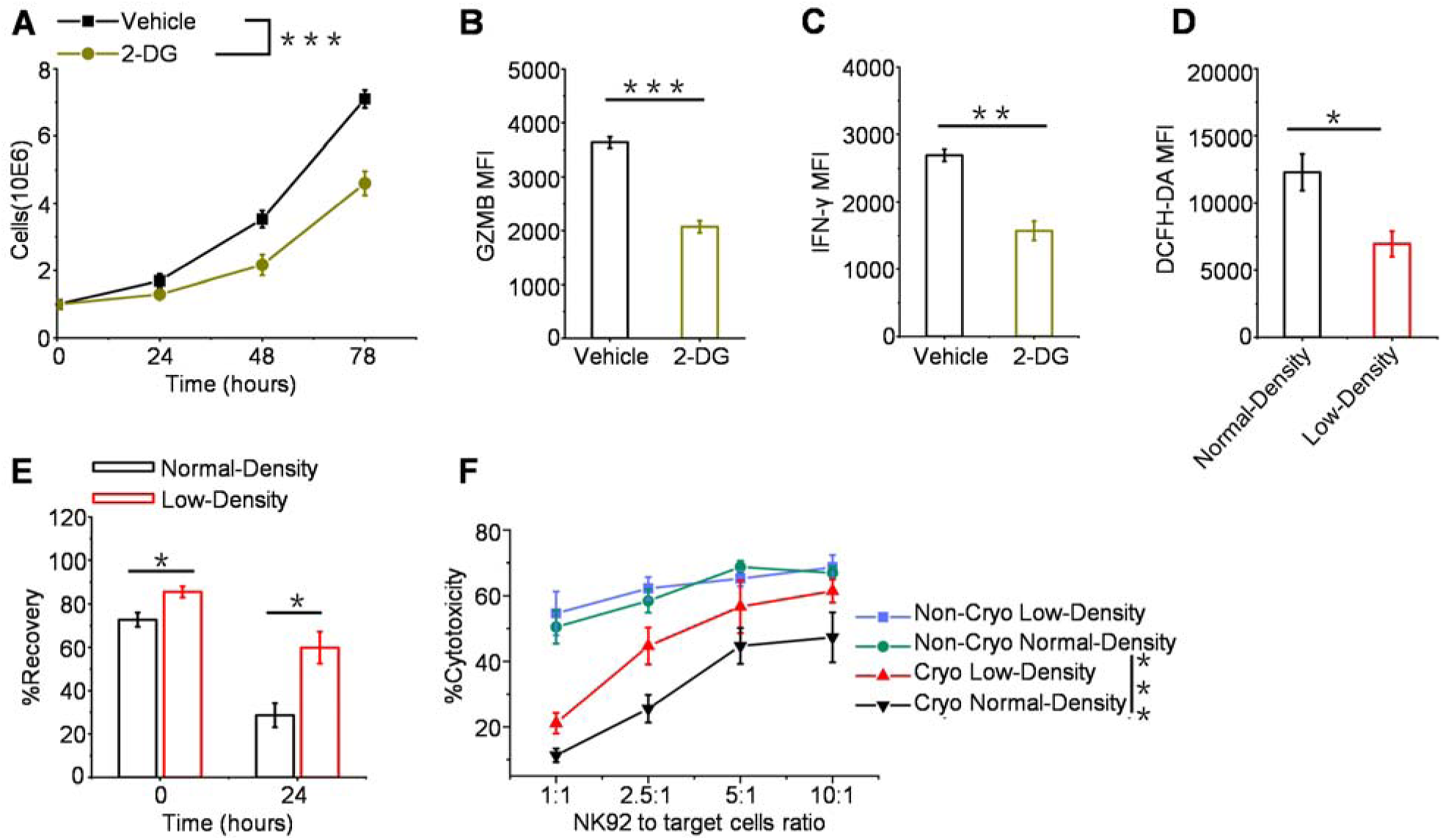
Glucose metabolism and ROS underlie NK92 cell cryo-vulnerability. (A) Proliferation of NK92 cells treated with 2-DG (5 mM). (B) GZMB levels of NK92 cells treated with 2-DG (5 mM). (C) IFN-γ levels of NK92 cells treated with 2-DG (5 mM). (D) ROS levels of NK92 cells cultured at different densities detected by DCFH-DA probes. (E) Recovery of NK92 cells cultured at different densities before cryopreservation, assessed at 0 and 24 hours post-thawing. (F) NK92 cytotoxicity assay with Raji cells. NK cells were cultured with Raji cells at the 1:1, 2.5:1 and 5:1 effector:target ratio for 4 h. Data are the mean ± SEM, n = 3 biologically independent experiments (A to F). Statistical analysis was performed using an unpaired two-tailed Student’s t test. Ns, P > 0.05, *P < 0.05, **P < 0.01, ***P < 0.001.

**Figure S4.**
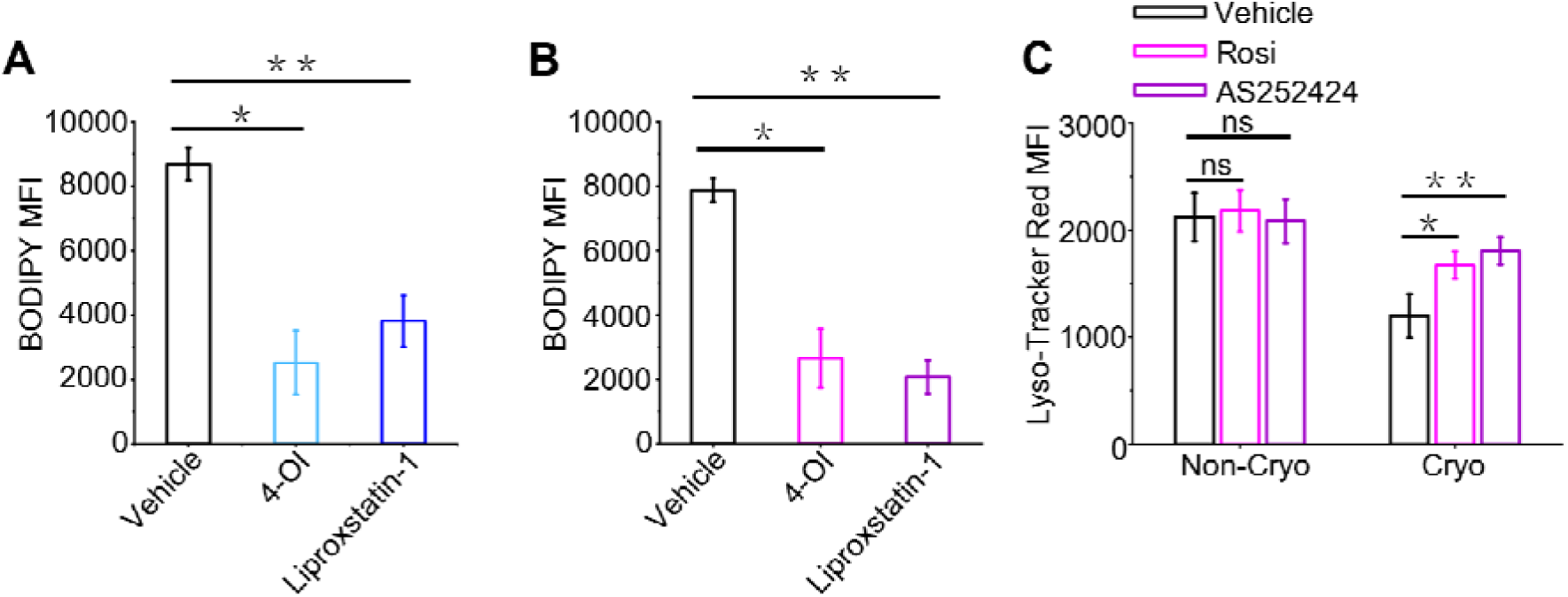
Effect of 4-OI or Liproxstatin-1 on lipid peroxidation of NK92 cells. (A) BODIPY oxidation in 4-OI (50 μM) or Liproxstatin-1 (10 μM)-treated NK92 cells. (B) BODIPY oxidation in 4-OI (50 μM) or Liproxstatin-1 (10 μM)-treated NK92 cells. (C) Lysosomal membrane permeability of Rosi (50 μM) or AS252424 (20 μM)-pretreated NK92 was detected by Lyso-Tracker Red probes. Data are the mean ± SEM, n = 3 biologically independent experiments. Statistical analysis was performed using an unpaired two-tailed Student’s t test. *P < 0.05, **P < 0.01.

**Figure S5.**
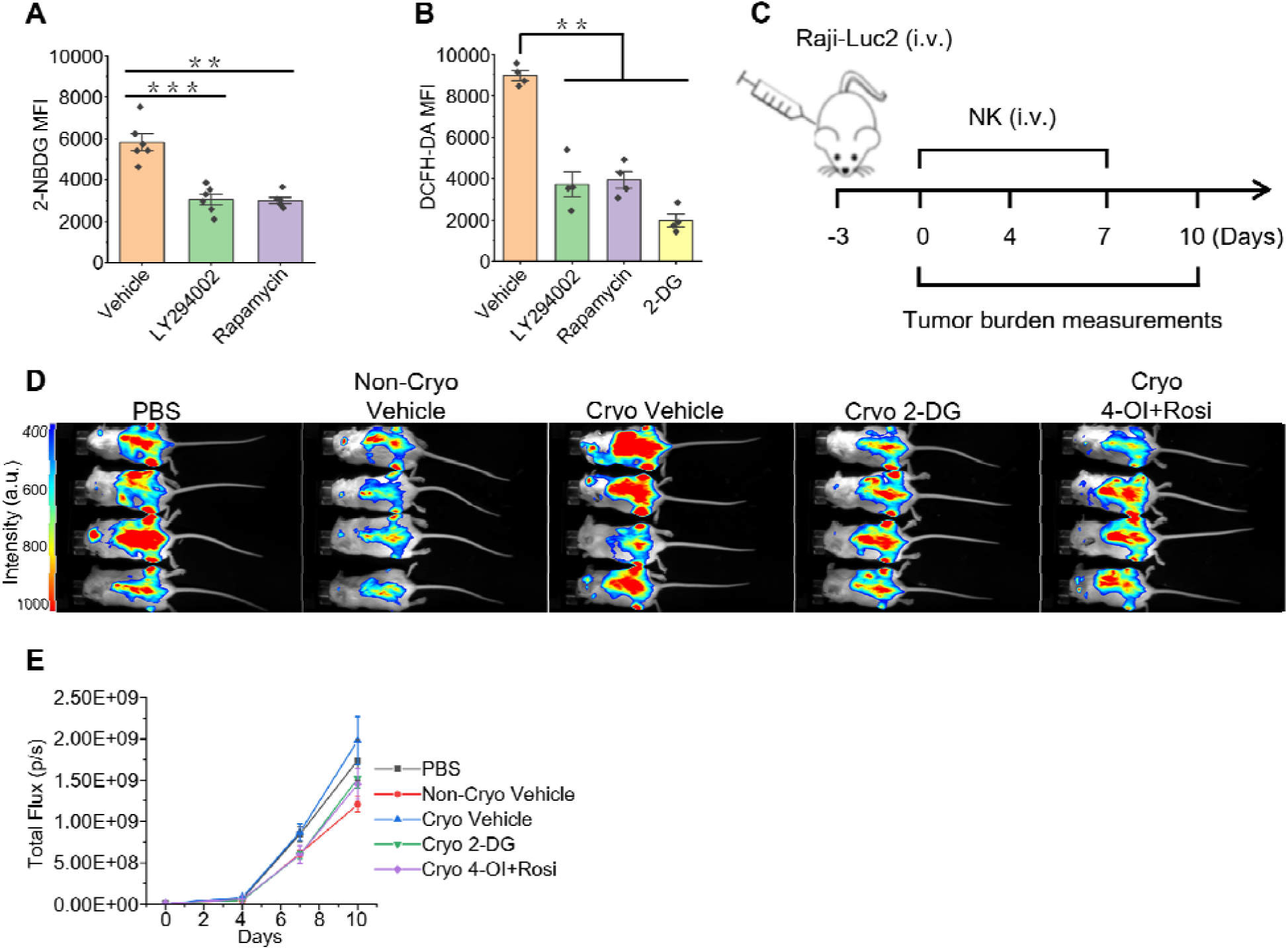
Mechanistic findings in NK92 cells translate to primary NK cells and improve antitumor efficacy in vivo. (A) 2-NBDG uptake by LY294002 (25 μM), Rapamycin (2 μM) or 2-DG (8 mM)-treated activated NK or T cells (n = 6 healthy donors). (B) ROS levels of LY294002 (25 μM), Rapamycin (2 μM) or 2-DG (8 mM)-treated activated NK or T cells detected by DCFH-DA probes (n = 4 healthy donors). (C) Schematic diagram for in vivo experiment. (D) Luminescent images of treated mice (n = 4 mice/group). (E) quantification of tumor burden (flux) (n = 4 mice/group). Data are the mean ± SEM (A, B, and E). Statistical analysis was performed using an unpaired two-tailed Student’s t test. **P < 0.01, ***P < 0.001.

**Figure S6.**
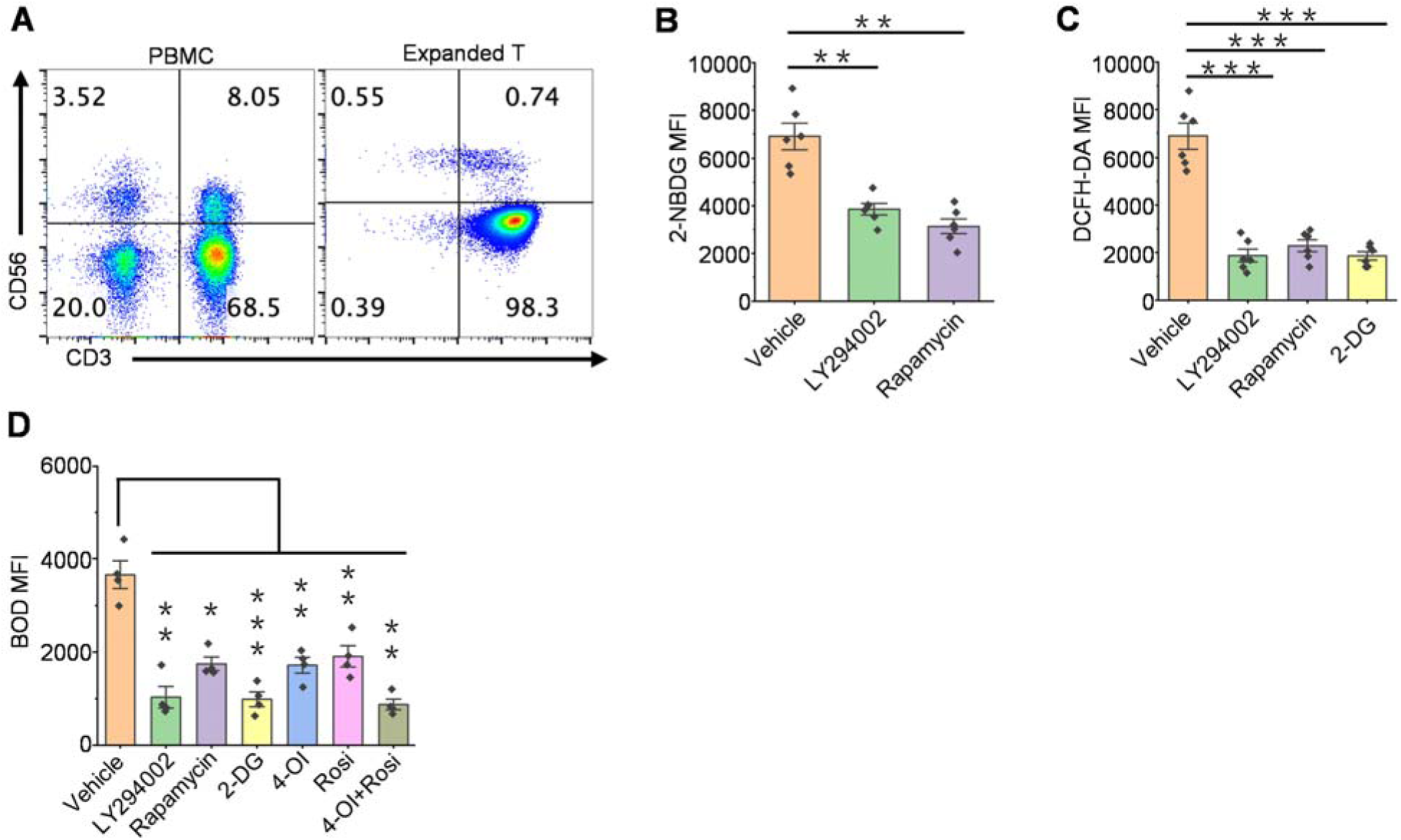
Generality of activation-induced vulnerability to cryopreservation. (A) Purity of peripheral blood derived αβ T cells (CD3^+^CD56^−^). (B) 2-NBDG uptake by LY294002 (25 μM), Rapamycin (2 μM) or 2-DG (8 mM)-treated activated αβ T cells (n = 6 healthy donors). (C) ROS levels of LY294002 (25 μM), Rapamycin (2 μM) or 2-DG (8 mM)-treated activated αβ T cells detected by DCFH-DA probes (n = 4 healthy donors). (D) BODIPY oxidation in LY294002 (25 μM), Rapamycin (2 μM), 2-DG (8 mM), 4-OI (50 μM) and/or Rosi (50 μM)-treated activated αβ T cells (n = 4 healthy donors). Data are the mean ± SEM (B to D). Statistical analysis was performed using an unpaired two-tailed Student’s t test. **P < 0.01, ***P < 0.001.

